# *Overdrive* is essential for targeted sperm elimination by *Segregation Distorter*

**DOI:** 10.1101/2024.06.04.597441

**Authors:** Jackson T. Ridges, Jackson Bladen, Thomas D. King, Nora C. Brown, Christopher R. L. Large, Jacob C. Cooper, Amanda J. Jones, Benjamin Loppin, Raphaëlle Dubruille, Nitin Phadnis

## Abstract

Intra-genomic conflict driven by selfish chromosomes is a powerful force that shapes the evolution of genomes and species. In the male germline, many selfish chromosomes bias transmission in their own favor by eliminating spermatids bearing the competing homologous chromosomes. However, the mechanisms of targeted gamete elimination remain mysterious. Here, we show that *Overdrive (Ovd)*, a gene required for both segregation distortion and male sterility in *Drosophila pseudoobscura* hybrids, is broadly conserved in Dipteran insects but dispensable for viability and fertility. In *D. melanogaster, Ovd* is required for targeted *Responder* spermatid elimination after the histone-to-protamine transition in the classical *Segregation Distorter* system. We propose that *Ovd* functions as a general spermatid quality checkpoint that is hijacked by independent selfish chromosomes to eliminate competing gametes.

---

Mendelian segregation is the foundation on which our understanding of genetics and population genetic theory rests. Segregation distorters are selfish chromosomes that violate Mendel’s Laws by over-representing themselves in the mature gamete pool of individuals that carry them (*1*, *2*). In the male germline, distorters act by selectively eliminating spermatozoa that carry their competing homologous chromosomes (*3*). Although a few segregation distorter genes have been identified, the mechanisms underlying the selective elimination of competing gametes remain poorly understood (*4–7*).

In *Drosophila*, some selfish chromosomes act by directly disrupting target chromosome segregation during meiosis (*5*). In other distorter systems, drive mechanisms involve the elimination of post-meiotic spermatids (*8–10*). During *Drosophila* spermiogenesis, groups of interconnected spermatids elongate and condense their nuclei into highly compact, needle-shaped sperm heads (*11*, *12*). This is achieved through a global chromatin remodeling process where almost all histones are progressively replaced first by transition proteins, such as Tpl94D, which are in turn replaced by Sperm Nuclear Basic Proteins (SNBPs), such as the Protamine-like ProtA and ProtB (*13*, *14*). One of the best-described distorter systems is *Segregation Distorter (SD)* in *Drosophila melanogaster* (*8*, *15*). In this system, selfish *SD* chromosomes selectively disrupt the histone-to-protamine transition in spermatid nuclei that carry an *SD*-sensitive homolog known as *Responder (Rsp)* (*16–18*). In *SD/Rsp* males, *Rsp* spermatid nuclei fail to condense properly and are eliminated during spermatid individualization. This is a common stage of spermiogenic failure observed in a variety of perturbations including male sterile mutants, high temperature induced sterility, etc. (*19*, *20*). This has led to speculation that a general male germline checkpoint may exist during spermatid individualization (*16*, *21*, *22*). However, the mechanisms of elimination of *Rsp* sperm remain unknown.

Intra-genomic conflict involving segregation distorters provides a leading explanation for the rapid evolution of hybrid male sterility but evidence for this idea remains scarce (*23–27*). A direct line of evidence connecting intra-genomic conflict to speciation comes from hybrids between two very young subspecies: *Drosophila pseudoobscura bogotana* (*Bogota*) and *Drosophila pseudoobscura pseudoobscura* (*USA*) (*28*). F1 hybrid males from crosses between *Bogota* mothers and *USA* fathers are nearly sterile but become weakly fertile when aged (*29*). These aged hybrid males produce nearly all female progeny due to segregation distortion by the *Bogota X*-chromosome. Although the genetic basis of segregation distortion and male sterility in these hybrids involves a complex genetic architecture, a single gene *Overdrive* (*Ovd*) is involved in both phenomena (*7*, *30*). Yet, little is known about the normal function of *Ovd* within species or its mechanistic role in segregation distortion and male sterility between species.

First, we wanted to understand the origins and patterns of molecular evolution of *Ovd*. To date the origins of *Ovd*, we used reciprocal BLAST and synteny analyses to search for orthologs of *Ovd* across *Diptera*. *Ovd* is present in all *Drosophilidae* and *Schizophora* species analyzed, but not detected outside of *Schizophora* (Figure 1A). Based on time-calibrated phylogenies of *Dipterans* and *Schizophora* (*31*, *32*), we conclude that *Ovd* originated at least 60 million years ago in the ancestor of *Schizophora* and has since been maintained without loss. Hybrid sterility and segregation distortion genes tend to change rapidly under recurrent positive selection (*5*, *7*, *33*, *34*). We used PAML (Phylogenetic Analysis by Maximum Likelihood) (*35*) to search for signatures of recurrent positive selection in *Ovd* in the *melanogaster* and *obscura* clades. Surprisingly, we did not observe high d_N_/d_s_ values for *Ovd*, nor did any individual lineage show accelerated accumulation of nonsynonymous substitutions (Figure 1B). Models of molecular evolution allowing for positive selection were not a significantly better fit than those that excluded it (*p =* 0.972 for *melanogaster* group species and *p =* 0.999 for *obscura* group species, likelihood ratio test). No individual residues within *Ovd* showed signatures of positive selection. Despite being involved in both segregation distortion and hybrid sterility, *Ovd* is surprisingly conserved and shows no signs of accelerated evolution.

**Figure 1:**
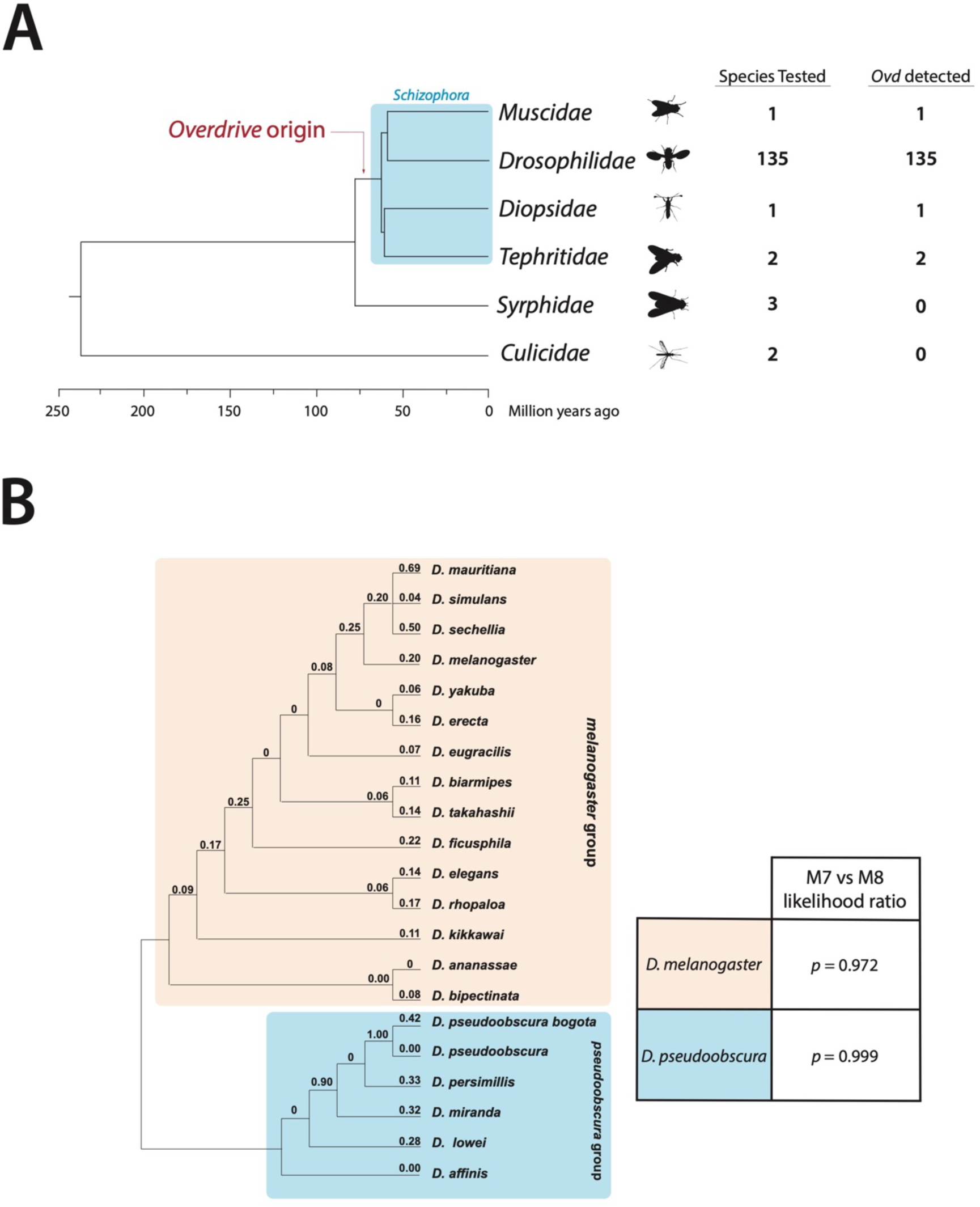
*Overdrive* has been maintained without loss since its origin at least 65 million years ago and does not evolve rapidly under positive selection. (A) We searched for orthologs of *Ovd* in 139 species of *Dipterans* using reciprocal BLAST and synteny checks. Orthologs of *Ovd* were detected in all *Schizophora* species tested but not detected outside of *Schizophora*. Time-calibrated phylogenies of *Schizophora* and Dipterans allow time estimates for our phylogeny (*31*, *32*). **(B)** Cladogram of *Ovd* sequence evolution across *melanogaster* (orange) and *pseudoobscura* (blue) groups with d_N_/d_S_ values for each branch calculated with the PAML package. *Ovd* does not evolve under recurrent positive selection as denoted by p-values from M7 vs M8 model comparisons.

Because *Ovd* is a conserved gene and can cause sterility and segregation distortion in *Bogota-USA* hybrids, we wondered if it is essential for male germline development. We used CRISPR/Cas9-based gene editing to generate null mutants of *Ovd* in both the *USA* and *Bogota* subspecies (Figure 2A). Surprisingly, *Ovd*-null individuals from either subspecies were fertile. We observed no difference in male fertility, sex-chromosome segregation ratios, or viability between *Ovd^Δ^* and wild-type controls in either subspecies (Figure 2B, 2C, 2D). *Ovd* thus appears to be dispensable for viability and male germline development in *D. pseudoobscura*.

**Figure 2:**
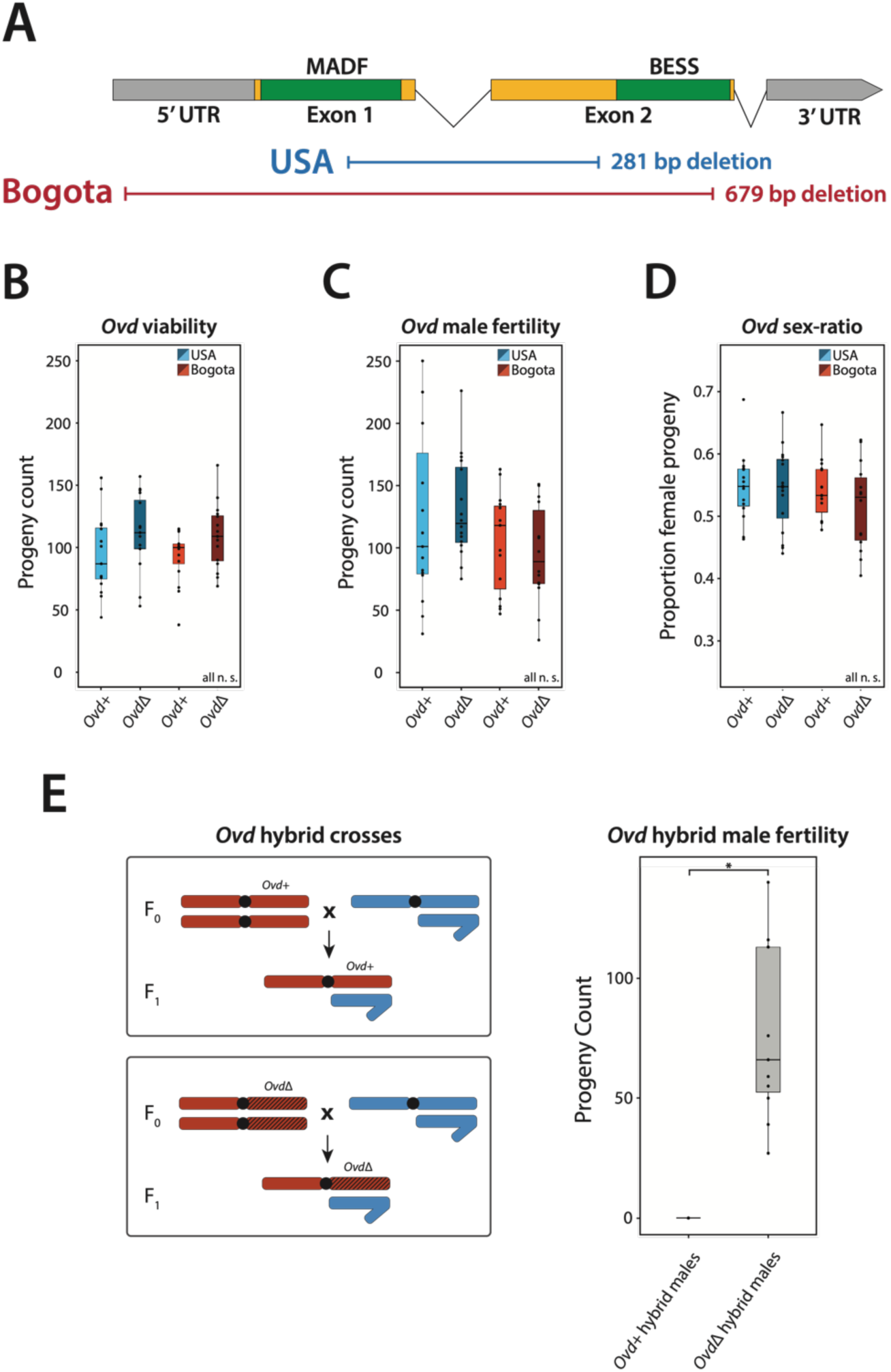
*Ovd* is non-essential in *D. pseudoobscura* in pure species but removing it restores fertility and normal segregation in interspecies F1 hybrid males. (A) Schematic showing CRISPR-Cas9 NHEJ knockouts of *Ovd* where nearly the whole coding region is deleted in *USA* and *Bogota* subspecies of *D. pseudoobscura*. **(B)** Viability of *Ovd^Δ^* and wild-type crosses in both *D. pseudoobscura* subspecies. Viability of *Ovd^Δ^* and wild-type individuals are not significantly different (*p*=0.10 for *USA*; *p*=0.056 for *Bogota*; two sample Student’s *t*-test) **(C)** Male fertility of *Ovd^Δ^* and wild-type individuals are not significantly different (*p*=0.73 for *USA*; *p*=0.54 for *Bogota*; two sample Student’s *t*-test). **(D)** Sex-ratio of progeny produced by *Ovd^Δ^* and wild-type males in both *D. pseudoobscura* subspecies. Progeny sex-ratios between *Ovd^Δ^* and wild-type individuals are not statistically significant (*p*=0.87 for *USA*; *p*=0.33 for *Bogota*; two sample Student’s *t*-test). **(E)** Crossing schematic and fertility of F1 hybrid males produced by crossing either *Ovd^Δ^ or Ovd^+^ Bogota* females to *USA* males. Deleting *Ovd* restores fertility to otherwise sterile F1 hybrid males (*p*=0.002; Wilcoxon test).

Previously, we showed that replacing an incompatible *Bogota* allele of *Ovd* with a compatible *USA* allele restores fertility and normal segregation to F1 hybrid males (*7*). However, it is not clear whether F1 hybrid males are sterile because they lack a *USA* allele of *Ovd* that provides an essential function or because the presence of the *Bogota* allele blocks male germline development. To discriminate between these possibilities, we crossed *Bogota* females with or without *Ovd* to *USA* males to produce F1 hybrid males (Figure 2E). In the control *Ovd^+^* cross, all F1 hybrid males were sterile, as expected. In contrast, *Ovd^Δ^* hybrid F1 males were fertile and showed normal sex-chromosome segregation ratios. We conclude that *Ovd* is not required for normal male germline development in pure species, but the presence of *Bogota Ovd* can block male germline development in hybrids.

Although *Ovd* is not an essential gene, its conservation across long evolutionary timescales suggests that it may have an important function. We hypothesized that *Ovd* may be involved in a putative male germline checkpoint. An *Ovd*-mediated checkpoint could explain three distinct observations. First, *Ovd* is dispensable for normal germline development because removing such a checkpoint would have no discernable effect in the absence of perturbation. For example, null mutants of known DNA damage checkpoint genes, such as *p53* and *Chk-2 kinase*, are viable and develop normally, but only show an effect in the presence of DNA damage (*36*, *37*). Second, removing *Ovd* restores normal segregation in hybrids. If an *Ovd*-mediated checkpoint specifically arrests only spermatids targeted for elimination, this would cause segregation distortion. Third, removing *Ovd* restores hybrid male fertility. If an *Ovd*-mediated checkpoint arrests all spermatids, this would cause sterility (Figure 3A).

**Figure 3:**
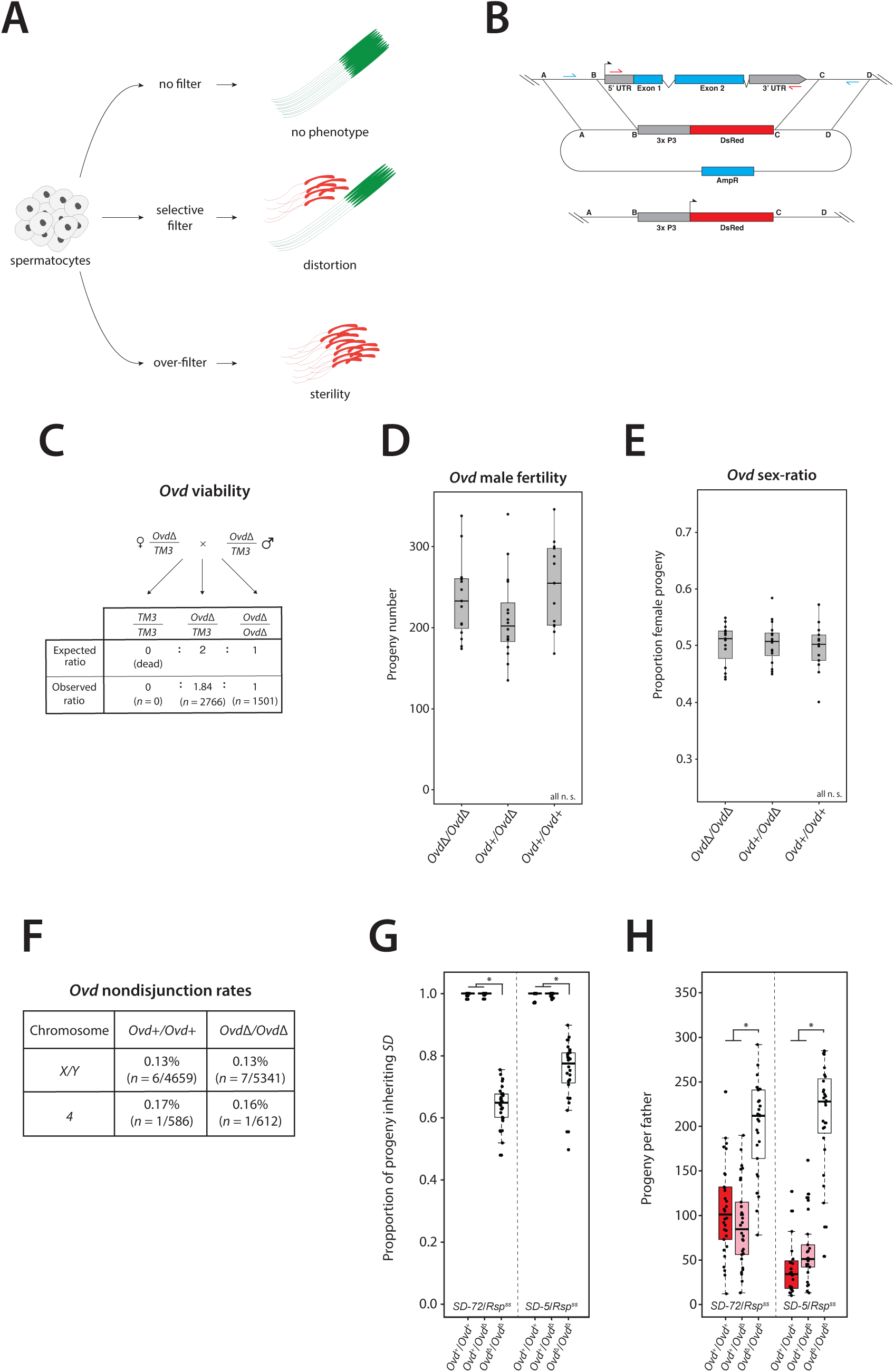
*Overdrive* is non-essential for viability and fertility in *D. melanogaster*. **(A)** A model for a role for *Ovd* in a germline checkpoint can explain why *Ovd* is dispensable for male germline development but is required for segregation distortion and male sterility. **(B)** Schematic for the design of *D. melanogaster Ovd* knockout. We use CRISPR-Cas9 to replace the coding sequence of *Ovd* with DsRed, a visible marker. **(C)** Progeny counts for *Ovd^Δ^* crosses show that it is not essential for viability. *Ovd^Δ^* shows no viability deficit relative to TM3 balancer. The number of progeny of *Ovd^Δ^* /TM3 heterozygote pairs did not deviate significantly from the expected Mendelian ratio of 2:1 *Ovd^Δ^*/TM3:*Ovd^Δ^*/*Ovd^Δ^* (p = 0.08, Cochran-Mantel-Haenszel test). **(D)** Male fertility for *Ovd^Δ^*. The number of progeny produced per male for wildtype *Ovd*, heterozygous for *Ovd^Δ^*, and homozygous for *Ovd^Δ^* are not significantly different from each other (one-way ANOVA; p = 0.124) **(E)** Sex-ratio of progeny produced by males wildtype for *Ovd*, heterozygous for *Ovd^Δ^*, and homozygous for *Ovd^Δ^* are not significantly differently from each other (one-way ANOVA; p = 0.875). **(F)** Nondisjunction rates for sex chromosomes and the fourth chromosome in *w^1118^* and *Ovd^Δ^*males. Sex chromosome nondisjunction frequencies do not differ significantly between the genotypes, nor do fourth chromosome nondisjunction frequencies. **(G)** *k*-values for *SD/Rsp^ss^* males with wild-type *Ovd*, heterozygous *Ovd^Δ^* and homozygous *Ovd^Δ^* for *SD-72* and *SD-5* chromosomes. Homozygous *Ovd^Δ^* males show reduced strength of distortion (*p* < 0.05, Three-way ANOVA, Tukey’s Honestly Significant Differences test). **(H)** Progeny count per male for *SD/Rsp^ss^* males with wild-type *Ovd*, heterozygous *Ovd^Δ^* and homozygous *Ovd^Δ^*. Homozygous *Ovd^Δ^* males produce more progeny than wild-type *Ovd* and heterozygous *Ovd^Δ^* males in an *SD* background (*p* < 0.05, Three-way ANOVA, Tukey’s Honestly Significant Differences test)

To test its hypothetical male germline checkpoint function, we switched to studying *Ovd* in *Drosophila melanogaster*, where more genetic and cytological tools are available. The predicted Ovd protein has a MADF DNA-binding domain and a BESS protein-interaction domain. MADF-BESS domain-containing proteins are involved in diverse processes such as transcriptional regulation and chromatin remodeling (*38–40*). We first studied the expression and localization pattern of Ovd in testes by introducing an N-terminus GFP-tag at the endogenous locus using CRISPR/Cas9. Testis imaging shows that Ovd is a nuclear protein expressed during spermatogenesis from male germline stem cells through the histone-to-protamine transition stages (Supplemental figure S1). Next, we used CRISPR/Cas9 to delete *Ovd* in *D. melanogaster* (Figure 3B, Supplemental figure S2). We observed little difference between male fertility, progeny sex-ratio, viability, or chromosomal non-disjunction rates between *Ovd^Δ^* and wild type controls (Figure 3C, 3D, 3E, 3F). Like in *D. pseudoobscura*, *Ovd* is not essential for normal male germline development in *D. melanogaster*.

If *Ovd* functions in a male germline checkpoint, then sperm that are normally eliminated in response to perturbations may develop to maturity when *Ovd* is removed. We turned to the *Segregation Distorter* (*SD*) system to perturb male germline development in *D. melanogaster*. We hypothesized that when *Ovd* is removed, *SD* chromosomes would fail to eliminate gametes bearing homologous chromosomes with *Rsp* satellite repeats. We first measured the strength of distortion by the *SD-72* chromosome against *Rsp^ss^*, a supersensitive *Rsp* chromosome, and observed near-complete distortion as expected (*k* > 0.99). We then measured the strength of distortion by the same *SD-72* chromosome against *Rsp^ss^* in an *Ovd^Δ^* homozygous background. We observed a strong reduction in the strength of distortion by *SD-72* in the absence of *Ovd* (*k* ∼0.65; Figure 3G). We repeated our measurements with an independently isolated *SD* chromosome, *SD-5* (*8*). Again, we found a strong reduction in the strength of distortion by *SD-5* in an *Ovd* homozygous null background (Figure 3H). *SD* thus requires *Ovd* for full-strength distortion. Together, our results show that *Ovd* is required for both sex-chromosome distortion in *D. pseudoobscura,* and autosomal distortion in *D. melanogaster*.

We then investigated spermiogenesis in *SD* males in the presence or absence of *Ovd*. In *SD-72/Rsp^ss^* males, spermiogenesis appears highly disturbed with many abnormal spermatid nuclei being eliminated, a typical phenotype of *SD* males (*16*). In contrast, spermiogenesis in *SD-72/Rsp^ss^* males appeared overall normal in an *Ovd^Δ^* homozygous background (Figure 4A). We then examined the histone-to-protamine transition in these males using *Tpl94D-RFP* and *ProtB-GFP* transgenes. As previously observed (*16*), about half spermatid nuclei showed a delay in protamine incorporation and elongation defects in *SD* males (Figure 4B). These nuclei fail to individualize and are eliminated before their release into the seminal vesicle, the sperm storage organ in males. In contrast, in an *Ovd^Δ^* homozygous background, nearly all spermatids incorporate protamine and elongate properly (Figure 4A). Removing *Ovd* thus appears to restore proper histone-to-protamine transition and individualization to *Rsp* sperm even in the presence of *SD*. To assess the quality of sperm nuclei produced by *SD* males that lack *Ovd*, we stained seminal vesicles with an antibody that recognizes double stranded DNA (dsDNA). We have previously shown that this antibody can stain improperly compacted chromatin in mature *Rsp* sperm nuclei but not properly condensed sperm nuclei which are refractory to immuno-staining (*16*). Remarkably, we observed a significant increase of anti-dsDNA positive sperm in seminal vesicles of *SD*/*Rsp^ss^* males lacking *Ovd* compared to *SD*/*Rsp^ss^* males with *Ovd* (30% decondensed vs. 4% decondensed) (Figure 4C, 4D). We conclude that, when Ovd is removed, *Rsp* spermatids differentiate to maturity despite chromatin condensation defects caused by *SD*. Because these sperm produce viable *Rsp^ss^* progeny, these condensation defects are not fatal. Taken together, these results indicate that *Ovd* does not itself cause condensation defects in mature sperm but is instead required for the selective elimination of improperly condensed *Rsp* spermatids during individualization.

**Figure 4:**
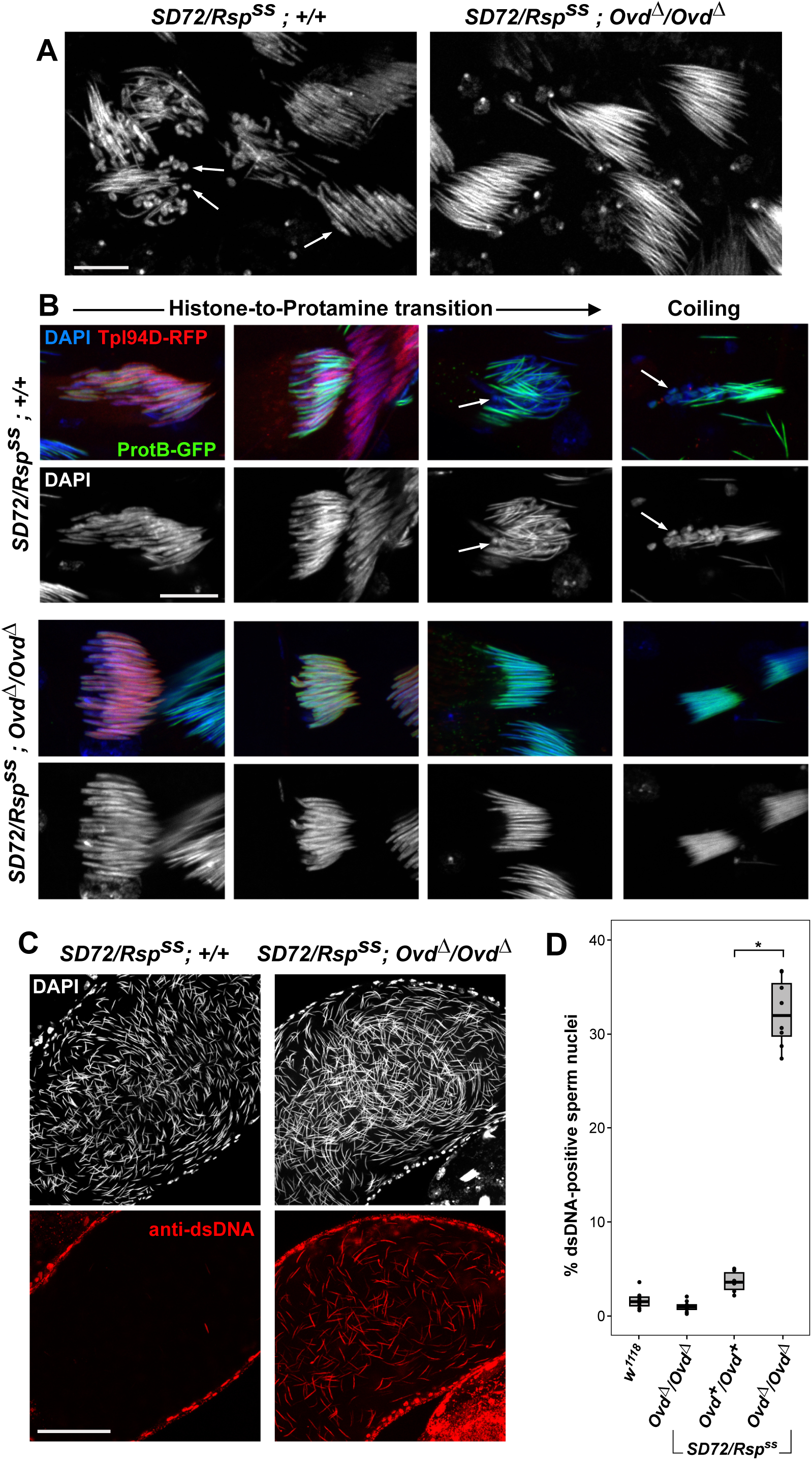
*Ovd* is required for selective elimination of *Rsp* sperm in the presence of *SD* in *D. melanogaster*. **(A)** Cysts of 64 spermatids in testes from *SD-72*/*Rsp^ss^* males with and without *Ovd* stained with DAPI. With Ovd, cysts are disorganized, half spermatid nuclei fail to elongate and degenerate (arrows). Without *Ovd*, spermatid elongation is restored. Bar: 10 µm (B) The histone-to-protamine transition of *SD-72*/*Rsp^ss^*males with and without *Ovd*. ProtB-GFP and Tpl94D-RFP transgenes are used to visualize protamines and transition proteins, respectively. With *Ovd*, half of the developing spermatids fail to incorporate protamines and do not elongate properly (arrows). Without *Ovd,* all sperm undergo the histone-to-protamine transition and elongation appropriately. Bar: 10 µm **(C)** Seminal vesicles of *SD-72*/*Rsp^ss^* males with and without *Ovd* stained with an anti-dsDNA antibody to reveal improperly condensed sperm nuclei. Bar: 50 µm. **(D)** Quantification of decondensed sperm in seminal vesicles of *SD-72*/*Rsp^ss^* males with and without *Ovd*, with *w^1118^*and *Ovd^Δ^*/*Ovd^Δ^* as controls. There is a significant increase in number of anti-dsDNA positive sperm nuclei from *SD-72/Rsp^ss^; Ovd^Δ^/Ovd^Δ^*males compared to *SD-72/Rsp^ss^; Ovd^+^/Ovd^+^*males (Wilcoxon Test; *p* < 0.01). Each dot corresponds to a male, decondensed sperm from one seminal vesicle was quantified per male.

*Ovd* is necessary for sex-chromosome segregation distortion and male sterility in *D. pseudoobscura* hybrids, and for autosomal segregation distortion in *D. melanogaster SD* males. Although *Ovd* is dispensable for normal spermiogenesis, it is evolutionarily conserved and blocks proper histone-to-protamine transition in *Rsp* spermatids in *D. melanogaster SD* males. Together, these properties suggest that the normal function of *Ovd* may involve the elimination of abnormal gametes during male germline development, which may get co-opted by selfish chromosomes.

It is worth noting the relationship between the genomic location of *Ovd* and its rate of evolutionary change. In *D. pseudoobscura, Ovd* is located on the distorting *Bogota X*-chromosome and rapidly accumulates non-synonymous changes in the *Bogota* lineage (*7*). In *D. melanogaster*, where *Ovd* is unlinked to the distorter, it is not rapidly evolving (*Ovd* is on the third chromosome, *SD* is on the second chromosome). *Ovd* thus evolves in a manner consistent with being locked in an evolutionary arms race in *D. pseudoobscura* but appears to act as an unwitting accomplice to *SD* in *D. melanogaster*. Nevertheless, *Ovd* is essential for two distant non-orthologous segregation distorters raising the possibility that independent selfish chromosomes may operate through shared mechanisms. Together, our findings open the door to understanding how mechanisms of gamete elimination may be involved in the evolution of selfish chromosomes within species and contribute to hybrid sterility between species.

## ACKNOWLEDGEMENTS

We thank Kent Golic for helpful discussions and feedback in improving this manuscript. We thank Titine Loppin for her continued support. Strains are available upon request. This work was supported by the National Institute of Health grant R01GM141422 to NP and French National Research Agency grant ANR-21-CE13-0037 to BL. We acknowledge the contribution of Lyon SFR Biosciences (UAR3444/CNRS, US8/INSERM, ENS de Lyon, UCBL) imaging facility (PLATIM) and fly food production (Arthrotools).

## SUPPLEMENTAL INFORMATION

### MATERIALS AND METHODS

#### Experimental Resources

##### Plasmids and Primers

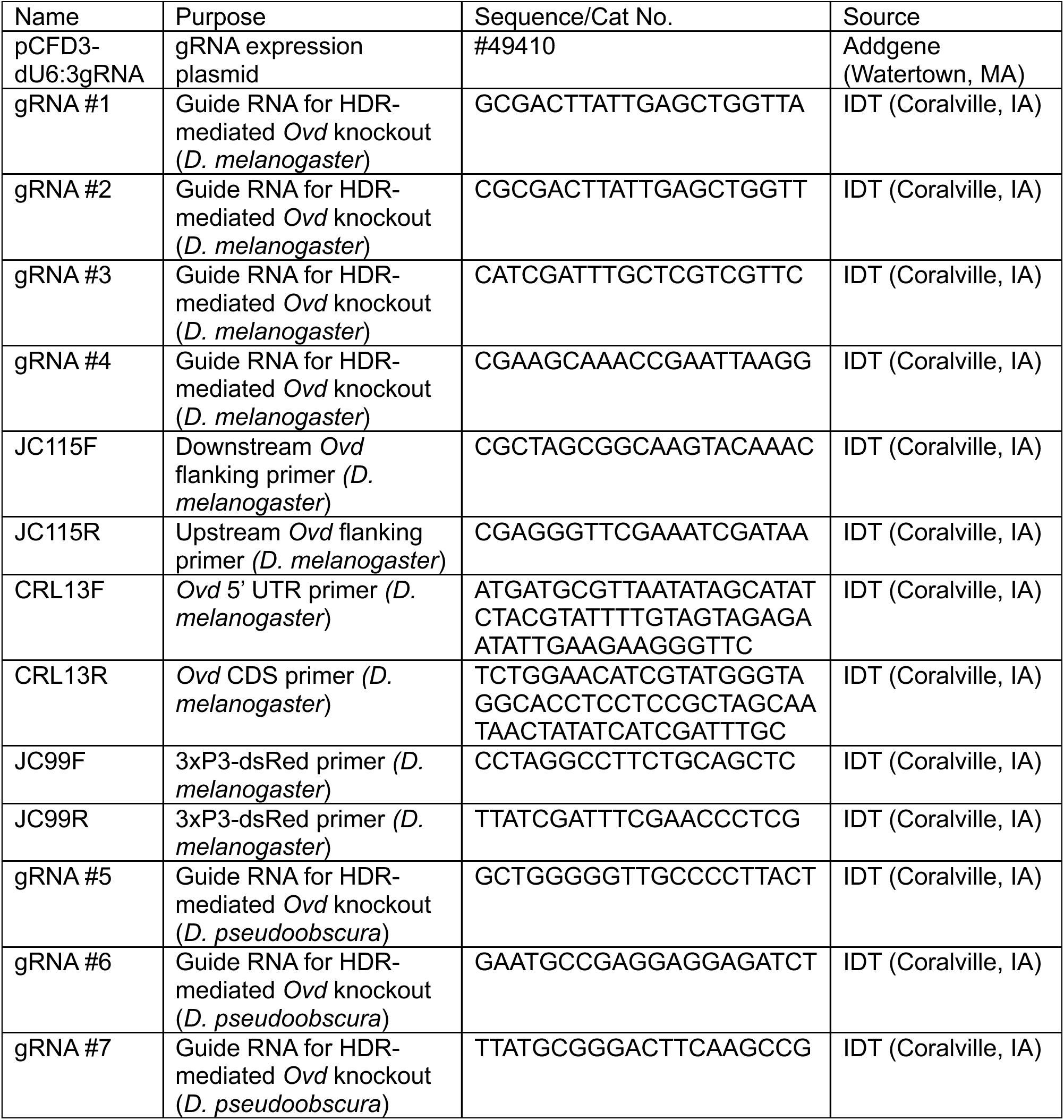
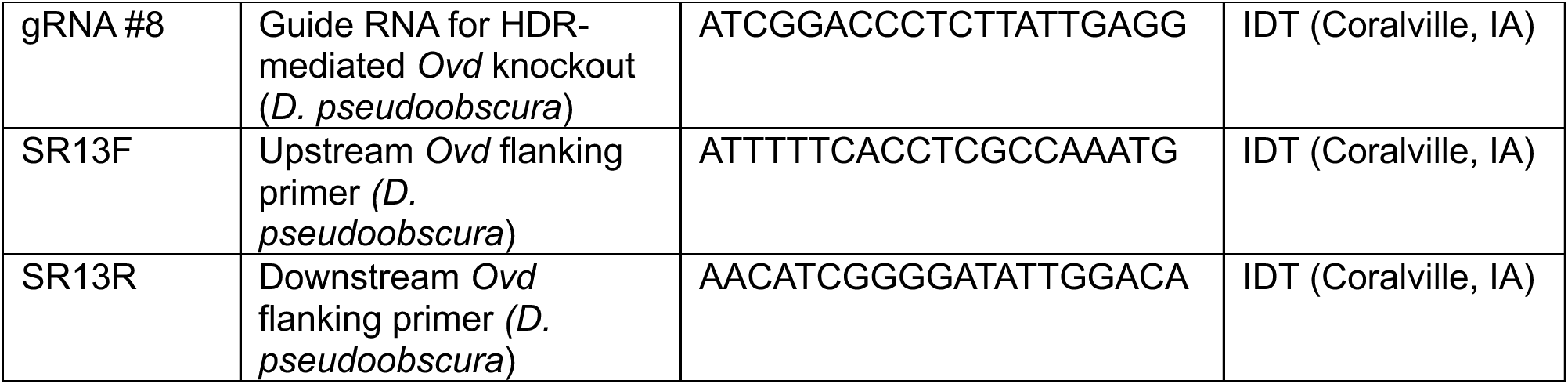

##### Fly Strains

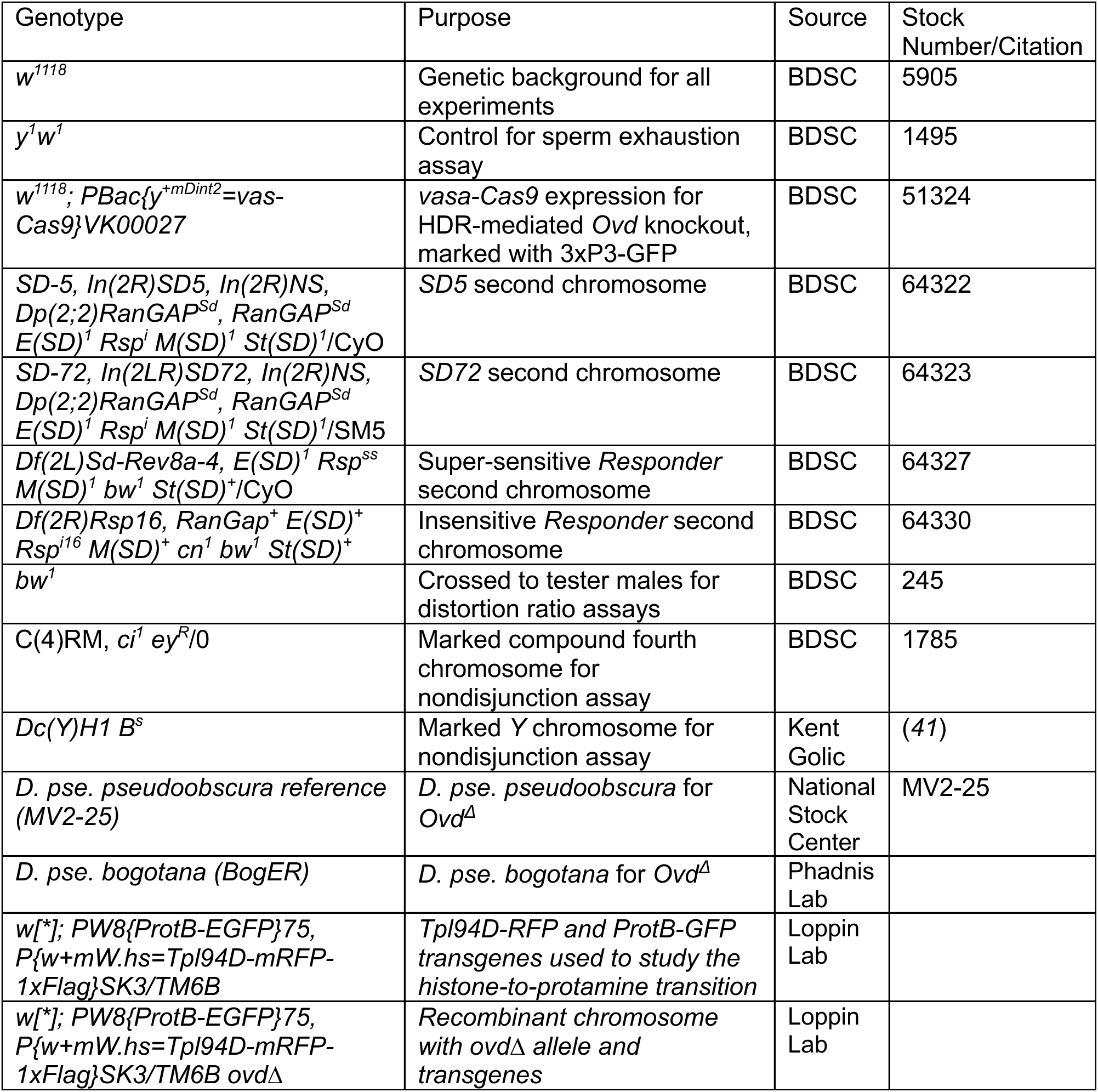

### CRISPR knockout of Overdrive in D. pseudoobscura

*Overdrive* knockout (*OvdΔ*) flies were generated using CRISPR/Cas9 based non-homologous end joining to create deletions of *Ovd* in *D. pseudoobscura pseudoobscura (USA)* and *D. pseudoobscura bogotana (Bogota).* First, gRNAs 5-8 (see plasmids and primers) were designed using CRISPR Optimal Target Finder (*42*). gRNAs were then complexed with Alt-R® CRISPR-Cas9 tracrRNA (IDT) and mixed with Alt-R® S.p. Cas9 Nuclease V3 (IDT). *USA* and *Bogota* embryos were injected with the complexed gRNAs and Cas9 nuclease.

To verify the loss of *Ovd,* primers flanking the native *Ovd* region were designed (SR13F and SR13R). Injected flies that produced short amplicons compared to wildtype controls were isolated. Sanger sequencing was done using the same primers to validate the deletion of *Ovd*. Flies with large deletions of the *Ovd* coding sequence were isolated in both subspecies (Figure 2A).

### Phenotyping D. pseudoobscura Overdrive knockout flies

For all *D. pseudoobscura* crosses, three males were crossed to three virgin females. Crosses were allowed to proceed for 7 days, then parents were removed. Progeny were counted at 28 days at 21 °C. Fifteen crosses were performed per genotype.

The viability of our *OvdΔ* mutations was assayed by comparing progeny counts of crosses with both parents homozygous for *OvdΔ* to crosses where both parents are wildtype. Two-sample Student’s t-tests were then used to analyze differences in progeny counts between wildtype and *OvdΔ* crosses for both subspecies, all differences were not significant. To assay male fertility and sex-ratio distortion, we crossed *OvdΔ* and wildtype males to wildtype females. Progeny counts and sex-ratio were compared between *OvdΔ* and wildtype crosses and differences were assessed with a two-sample Student’s t-test, all differences were not significant.

To generate *OvdΔ Bogota-USA* F1 hybrid males, homozygous *OvdΔ Bogota* females were crossed to *USA* males. F1 males were then crossed to *USA* females, and progeny were counted. Differences in fertility between *OvdΔ* F1 hybrid males and control wildtype F1 hybrid males were analyzed with a Pairwise Wilcoxon Rank Sum test.

### CRISPR knockout of Overdrive in D. melanogaster

*Overdrive* knockout (*OvdΔ*) flies were generated using CRISPR homology-directed repair to replace *Overdrive* with 3×P3-dsRed. Knockouts were made in a *vasa-Cas9* background (*w^1118^; PBac{y^+mDint2^=vas-Cas9}VK00027*). *Vasa-Cas9* embryos were injected with four pCFD3-dU6:3gRNA (Addgene, Plasmid #49410) vectors, each containing a gRNA with homology within the *Overdrive* coding sequence (*Plasmids and Primers* table). The donor template with 3×P3-dsRed2 was flanked with ∼1000 bp homology arms (Fig. 3A).

To verify the loss of *Ovd*, we used multiple primer pairs targeting *Ovd* and the region flanking the insertion site (*Plasmids and Primers* table; Fig. 3B). The first primer pair (JC115F and JC115R,) flanked *Ovd*, with the primers targeting approximately 2 kb upstream of the 5’ UTR and downstream of the 3’ UTR. These amplified in both *OvdΔ* and control *w^1118^* flies, showing a slightly larger product size in *OvdΔ* consistent with successful replacement with 3×P3-dsRed (expected product sizes 5577 bp with native *Ovd*, 6298 bp after allele swap with 3×P3-dsRed, Figure 3B). The second primer pair (CRL13F and CRL13R) respectively targeted the *Ovd* 5’UTR (approximately 450 bp upstream of the start codon) and *Ovd* coding sequence (immediately upstream of the stop codon). This primer pair produced amplicons in control *w^1118^*flies but not in *OvdΔ* flies, indicating loss of *Ovd* sequence in the knockout flies (Figure 3B). To verify that integration of our 3×P3-dsRed2 reporter construct occurred at the native *Ovd* locus, we used a third primer pair (JC99F and JC99R) with homology within the *dsRed2* coding sequence which only amplified in *OvdΔ* flies. Sanger sequencing reads from the *OvdΔ* PCR products confirmed that dsRed successfully replaced *Ovd* in our *OvdΔ* flies.

### Phenotyping *D. melanogaster Overdrive* knockout flies

*Viability Assay*: To assess the effects of *Overdrive* knockout on viability, we compared the viability of homozygous *OvdΔ* flies and *OvdΔ/TM3, Sb* flies. Eight *OvdΔ/TM3, Sb* males aged 0-3 days were mated on Day 1 with ten virgin females of the same genotype aged 0-3 days. This cross was conducted in 10 biological replicates. Parents were allowed to lay in the original vial for Days 1-4, and transferred to new vials every three days for a total of three times. Progeny in each vial were counted 18 days after the parents were introduced. A Cochran-Mantel-Haenszel test was applied to determine if the progeny deviated from the expected distribution of 2:1 *OvdΔ/TM3, Sb* heterozygotes to *OvdΔ* homozygotes.

*Sex Ratio Assay:* Although *Overdrive* appears to function as a selfish element in the *D. pseudoobscura* system that eliminates Y-bearing sperm to drive its own transmission, we wanted to exclude the possibility that *Overdrive* perturbation instead has a general male-killing effect. We therefore compared the sex ratio of the progeny of *OvdΔ* homozygote fathers and control *w^1118^* fathers. Single male flies aged 1-2 days were crossed to three *w^1118^* virgins aged 0-3 days. Flies were mated for one day, transferred to a new vial, and allowed to lay for three days. Parents were then cleared from the vial. Progeny were counted 14 days after parents were cleared. Total numbers of male and female progeny for each genotype were summed across biological replicates, and differences in male and female progeny numbers between genotypes were evaluated using the Student’s two-sample *t*-test.

*Nondisjunction Assays:* For all nondisjunction assays, individual males aged 1-2 days (30 males total per genotype) were crossed with three virgins aged 0-3 days (female genotype was *w^1118^* for sex chromosome assay; *C*(*4*)*RM* for fourth chromosome assay). Parents were transferred into a new vial after three days and cleared after six days. Progeny were scored and counted on Day 18.

We compared nondisjunction rates between *OvdΔ* fathers and *w^1118^* fathers for the sex chromosomes and the fourth chromosome. To measure sex chromosome nondisjunction, we used the *DcY(H1*) chromosome, which contains a translocation of the dominant eye marker *B^S^*onto the *Y* chromosome. *H1* chromosomes were introduced into wild-type (*w^1118^*) and *OvdΔ*/*OvdΔ* backgrounds. This allowed us to distinguish offspring resulting from nondisjunction events (*XO* males with wild-type eyes and *XXY* females with *Bar* eyes) from normal progeny (*XY* males with *Bar* eyes and *XX* females with wild-type eyes).

For fourth chromosome nondisjunction assays, we mated *OvdΔ* and *w^1118^* males with compound-fourth females (*C*(*4*)*RM*, *ci^1^ ey^R^*/0). Viable haplo-4 and triplo-4 progeny are produced by normal haplo-4 sperm fertilizing nullo-4 or compound-4 eggs. If a nullo-4 sperm resulting from a male nondisjunction event fertilizes a compound-4 egg, the progeny are recognizable due the recessive markers on the maternal compound chromosome. Nullo-4 zygotes are inviable, and while tetra-4 progeny can also arise from male nondisjunction events, they are undetectable in this assay (*43*). Fourth chromosome nondisjunction assays were repeated with *OvdΔ* on a pure *w^1118^* background and on a *Rsp^i^*/CyO background, *Rsp^i^* being a *Rsp* chromosome insensitive to SD.

For sex chromosome assays, differences between the genotypes in the number of nondisjunction events were assessed with a two-sample Student’s t-test. For fourth chromosome assays, results were analyzed with pairwise two-sample *c*-tests (using the Poisson distribution) with the Bonferroni false-discovery correction.

*Cell Cycle Progression Assay:* We tested whether *Overdrive* knockout affected the response of somatic cells to radiation-induced double-strand breaks. Third instar *w^1118^* and *OvdΔ* larvae (aged approx. 4 days) were transferred to a laying plate and *X*-ray irradiated for 4,000 rads. After 30 minutes of recovery, wing and eye disc tissues were dissected in PBS. Tissues were then fixed for 1 hr in 4% paraformaldehyde solution (Life Biotechnologies, Carlsbad, CA), washed 2×15 minutes in PBS with 0.1% (w/v) sodium deoxycholate and 0.1% (v/v) Triton, and 2×15 minutes in PBX (PBS w/ 0.1% (v/v) Triton). Tissues were incubated overnight at 4°C with rabbit anti-His3S10Phos antibody (abcam, Cambridge, UK), washed 3×15 minutes in PBX, incubated in goat anti-rabbit Alexa 568 secondary (Life Biotechnologies, Carlsbad, CA), washed 3×15 minutes in PBX, stained with 0.1% DAPI, mounted in Vectashield, and sealed with nail polish. Z-stacks of tissues were imaged using a Zeiss LSM 880 Airy Scan confocal microscope. Maximum-intensity Z-projections were generated in Fiji, and the number of phospho-Histone3S10 foci were scored in discs of irradiated and non-irradiated larvae using the CellCounter plugin (*44*). Differences between the irradiated and non-irradiated treatments, and the *OvdΔ* larvae and *w^1118^* controls, were analyzed using Mann-Whitney tests.

### *Overdrive* Molecular Evolution

*Drosophila* species used for domain structure predictions and PAML: *melanogaster, simulans, sechellia, mauritiana, yakuba, erecta, eugracilis, biarmipes, takahashii, ficusphila, elegans, rhopaloa, kikkawai, ananassae, bipectinata* (*melanogaster* group) and *pseudoobscura pseudoobscura, pseudoobscura bogotana, persimilis, miranda, lowei,* and *affinis* (*obscura* group). All analyses were conducted using the reference genome assembly or whole-genome sequencing contig databases publicly available through NCBI.

*Ovd* domain structure: We aligned orthologous *Ovd* sequences across the *melanogaster* and *obscura* groups, compared their predicted secondary structures, and analyzed their rates of nucleotide substitution. *Overdrive* orthologs from 15 *melanogaster* group species and six *obscura* group species (see *Fly Strains* section for list of species) were identified using reciprocal best-hits BLAST (tBLASTn and BLASTx). We then aligned these sequences using default settings in the MUSCLE alignment program and generated visualizations of conservation rates using Jalview (*45*, *46*). To investigate the degree of secondary structure conservation between the *D. pseudoobscura* and *D. melanogaster* orthologs of *Overdrive*, we predicted the location of alpha helices and beta sheets for the two *Ovd* orthologs and the analogous MADF– and BESS-containing proteins *Adf1* and *Dlip3* using the JNetPRED, JNetHMM, and JNetPSSM models in the JPred software package (*47*).

*PAML*: Analysis of *Overdrive* sequence evolution was conducted according to the method described in Cooper and Phadnis (*48*). Because we hypothesized that *Overdrive* could display different evolutionary dynamics in the *melanogaster* and *obscura* groups, and to avoid the possibility of synonymous site saturation, we analyzed the 15 *melanogaster* group species and six *obscura* group species separately. For each group, we used the Phlyogenetic Analysis by Maximum Likelihood (PAML) package (*35*). We first used the codeml feature in PAML to estimate d_N_/d_S_ for each branch in the gene tree of each species group. To test for positive selection within each species group, we compared NS sites models M7 (which allows only for purifying selection and neutral evolution) and M8 (which allows a class of sites within the sequence to have d_N_/d_S_ values greater than 1). In the results, we present the *p*-value of the likelihood-ratio test using the log-likelihood scores for the two models. A *p*-value under 0.025 (after Bonferroni correction for multiple comparisons) would indicate that the M8 model was a better fit for the sequence data, suggesting a pattern of recurrent positive selection within the coding sequence.

*Ovd ortholog search*: The following *Drosophilidae* species genome assemblies were used for ortholog searches of *Ovd*:

*Leucophenga varia, Scaptodrosophila lebanonensis, Chymomyza costata, Lordiphosa mommai, Lordiphosa magnipectinata, Lordiphosa clarofinis, Lordiphosa stackelbergi, Lordiphosa collinella, Drosophila sturtevanti, D. neocordata, D. saltans, D. prosaltans, D. sucinea, D. nebulosa, D. insularis, D. tropicalis, D. willistoni, D. equinozialis, D. paulistorum, D. subobscura, D. guanche, D. bifasciata, D. obscura, D. tristis, D. ambigua, D. lowei, D. miranda, D. pseudoobscura pseudoobscura, D. pseudoobscura bogotana, D. persimilis, D. azteca, D. Athabasca, D. varians, D. ercepeae, D. pseudoananassae, D. malerkotiana, D. parabipectinata, D. bipectinate, D. ananassae, D. oshimai, D. elegans, D. fuyamai, D. kurseongensis, D. carrolli, D. rhopoloa, D. ficusphila, D. biarmipes, D. subpulchrella, D. suzukii, D. takahashii, D. eugracilis, D. melanogaster, D. mauritiana, D. sechellia, D. simulans, D. orena, D. erecta, D. teissieri, D. yakuba, D. santomea, D. pectinifera, D. trapezifrons, D. triauraria, D. Auraria, D. tani, D. rufa, D. lacteicornis, D. asahinai, D. kanapiae, D. kikkawai, D. leontia, D. bocki, D. watanabei, D. punjabiensis, D. truncate, D. mayri, D. jambulina, D. seguyi, D. vulcana, D. bakoue, D. tsacasi, D. nikananu, D. burlai, D. bocqueti, D. busckii, D. repletoides, Zaprionus inermis, Zaprionus kolokinae, Zaprionus tsacasi, Zaprionus ornatus, Zaprionus africanus, Zaprionus indianus, Zaprionus gabonicus, Zaprionus capensis, Zaprionus taronus, Zaprionus davidi, Zaprionus vittiger, Zaprionus lachaisei, Zaprionus nigranus, Zaprionus camerounensis, D. quadrilineata, D. pruinosa, D. immigrans, D. sulfurigaster, D. kepylauana, D. albomicans, D. nasuta, D. cardini, D. dunni, D. arawakana, D. funebris, D. innubila, D. lacertosa, D. robusta, D. melanica, D. littoralis, D. virilis, D. americana, D. novamexicana, D. nannoptera, D. pachea, D. hydei, D. navojoa, D. arizonae, D. mojavensis, Scaptomyza hsui, Scaptomyza polygonia, Scaptomyza graminum, Scaptomyza montana, Scaptomyza flava, Scaptomyza pallida, D. grimshawi, D. murphyi, D. sproati*

The following non-*Drosophilidae Dipterans* species genome assemblies were used for ortholog searches of *Ovd*:

*Aedes albopictus, Aedes aegypti, Volucella bombylans, Syritta pipiens, Xylota sylvarum, Anastrepha ludens, Ceratitis capitata, Teleopsis dalmanni, Musca domestica*

When multiple genome assemblies were available, we used NCBI reference genomes. For species without a reference genome, the most recent genome assembly on NCBI genomes as of January 1^st^, 2024 was used. To identify orthologs of *Ovd*, tblastn and reciprocal blastx were performed against each genome listed above. tblastn was done with the following parameters: – gapopen 11 –gapextend 1 –evalue .001. *D. melanogaster Ovd (CG6683)* was used as a query. When the reciprocal best hit was *D. melanogaster Ovd,* the hit was called as an ortholog. If no reciprocal best hit was called, synteny checks were performed to validate the loss of *Ovd*. Absence of *Ovd* was concluded when both reciprocal best hit and synteny checks indicated that *Ovd* was not present.

### Transmission Distortion and Fertility in an *SD* Background

To determine whether *Overdrive* knockouts alter the level of transmission distortion caused by *Segregation Distorter* (*SD*) chromosomes, we generated 18 different strains of tester males. These represented every possible combination of three driving (or non-driving) chromosomes (*SD-5*, *SD-72*, and a *CyO* control), two chromosomes targeted (or not) by drive (*Rsp^ss^* and *Rsp^i^*), and three *Overdrive* states (wild-type, homozygous knockout, and heterozygotes). We used two different driving chromosomes isolated from natural populations, *SD-5* and *SD-72*. In males, these *SD* chromosomes drive against Responder-sensitive or – supersensitive chromosomes. While all *SD* chromosomes carry the same *RanGAP* duplication, their pattern of rearrangements, strength of drive, and viability relative to wild-type vary, so we used two different *SD* chromosomes in our experiments. As a control, we used the CyO balancer chromosome, a second chromosome that is inverted and homozygous lethal like the *SD* chromosomes we used (*49*) but has no driving properties and is insensitive to drive.

We used two different chromosomes as targets for drive. The first line we used, *Rsp^ss^*, has a high copy number of *Responder* repeats, rendering it supersensitive to *SD*. The second line, *Rsp^i^*, has a deletion of the *Rsp* locus, making the chromosome insensitive to SD (See *Fly Strains* table for full genotypes). Finally, we tested three different *Overdrive* conditions: wild-type, homozygous deletion of *Overdrive* (*OvdΔ*/*OvdΔ*), and heterozygotes (*OvdΔ*/+).

We tested all possible combinations of *SD*, *Rsp*, and *Ovd* alleles. To quantify segregation distortion, we crossed the males to *bw^1^* virgins. In each cross, 30 males aged 1-7 days were placed singly in vials and each crossed to 3 virgins aged 4-10 days. Parents were flipped into new vials on Day 3 of the cross, and cleared on Day 6. Progeny were counted on Day 18. Since both the *Rsp* chromosomes we used were marked with *bw^1^*, we were able to determine that brown-eyed progeny had received a *Rsp* chromosome from their fathers. Quantifying the proportion of progeny with wild-type eyes thus gave us a measure of *k*, the coefficient of transmission distortion. Differences in *k* among the treatment groups were modeled with a three-way ANOVA (k ∼ Overdrive * Distorter * Responder) on arcsine-transformed data (since many of the values of *k* were near or equal to 1) and analyzed with Tukey’s Honestly Significant Differences test.

### Immunofluorescence and imaging

Immunostainings in whole mount testes and seminal vesicles were performed as previously described. Briefly, male gonads we dissected in 1X PBS-0.15% Triton (PBS-T), then fixed for 20min at RT in 4% formaldehyde in PBS-T. Then tissues were wash 3 times and mounted in mounting medium containing DAPI for direct observation of DAPI staining and/or native GFP and RFP fluorescence. For antibody stainings, after fixation and washes, gonads were incubated in primary antibody [1:3000 for the mouse anti-dsDNA antibody (Abcam ref# ab27156)] diluted in PBS-T overnight at 4°C. The next day, tissues were washed 3 times in PBS-T and incubated with secondary antibody (1:500 Jackson Immunoresearch) at RT for 2-3 hrs. They were then mounted in mounting medium containing DAPI (2µg/mL). Testes and seminal vesicles were imaged with a LSM800 confocal microscope (CarlZeiss) and images were processed with Fiji software.

**Supplemental Figure S1:**
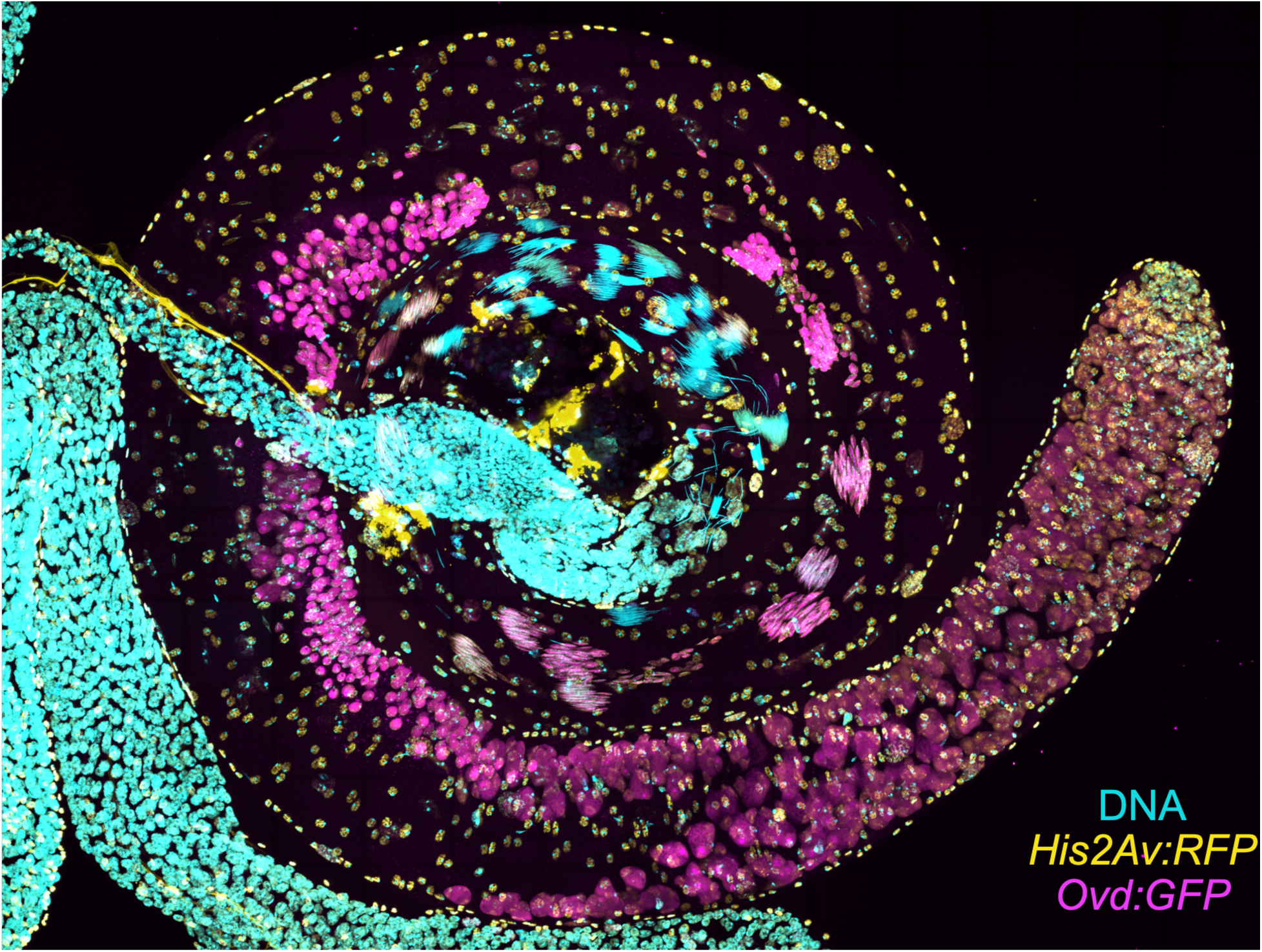
*Overdrive* is expressed in the male germline of *D. melanogaster* from male germline stem cells to canoe stage spermatids undergoing the histone-to-protamine transition. DNA is shown in cyan (DAPI), Histone2Av-RFP is shown in yellow, Ovd-GFP in magenta.

**Supplemental Figure S2:**
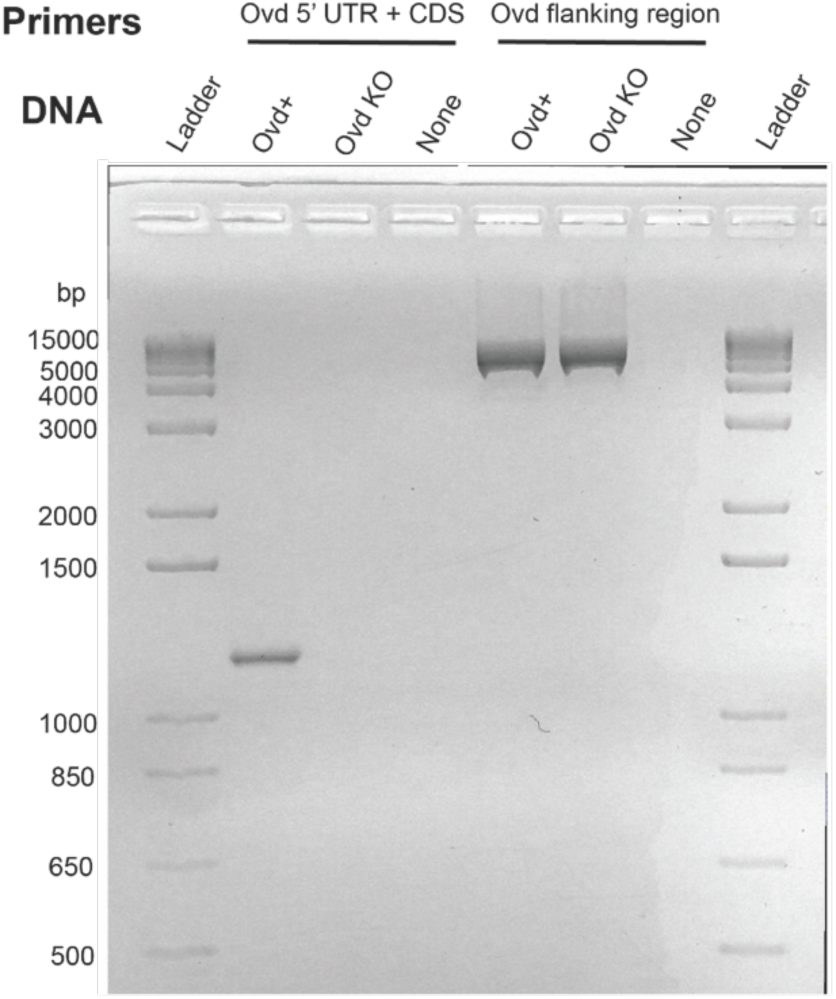
Validation of replacement of *Ovd* with DsRed. Primers placed inside of *Ovd* no longer amplify in *Ovd* KO, but primers flanking the gene still amplify.

**Supplemental Figure S3:**
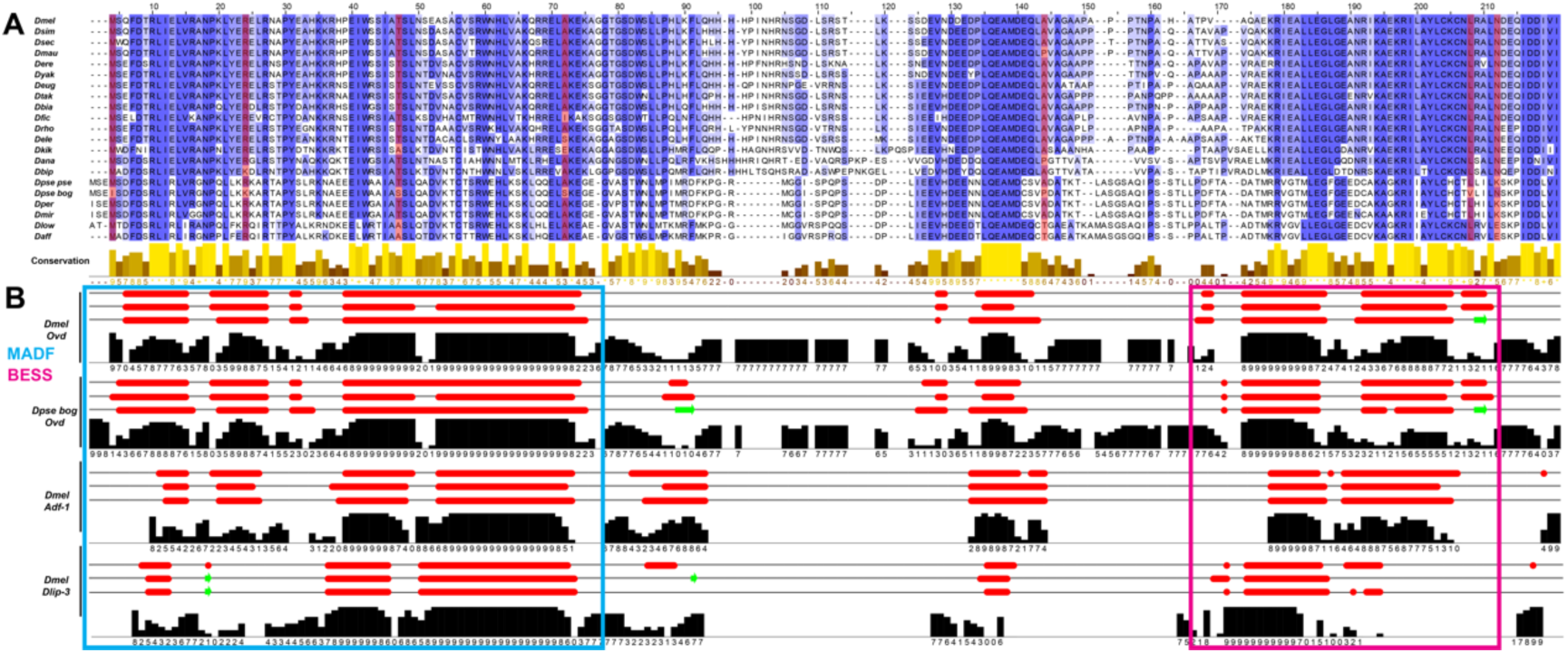
(A) Alignment of *Overdrive* coding sequence for 15 *melanogaster* and 5 *obscura* group species. Darker blue shading signifies a higher degree of amino acid sequence conservation. Yellow bars under the sequences represent the degree of conservation at each residue on a 0-9 scale (* = perfect conservation). Fixed amino acid differences in *D. pse. bogotana* (relative to *D. pse.* pseudoobscura) are shaded in red. **(B)** Predicted secondary structure for *Overdrive* in *D. pse. bogotana* and *D. melanogaster* based on the JNetPRED (top), JNetHMM (middle), and JNetPSSM (bottom) models in the Jpred package. Alpha helices are shown in red and beta sheets are shown in green. Histogram denotes confidence in the secondary structure calls for each residue on a 0-9 scale. Two proteins with MADF DNA-binding domains and BESS protein-protein interaction domains, *Adf1* and *Dlip3*, are shown for comparison. The cyan box denotes the approximate location of the MADF domain in all four proteins, and the magenta box denotes the approximate location of the BESS domain.

**Supplemental Figure S4:**
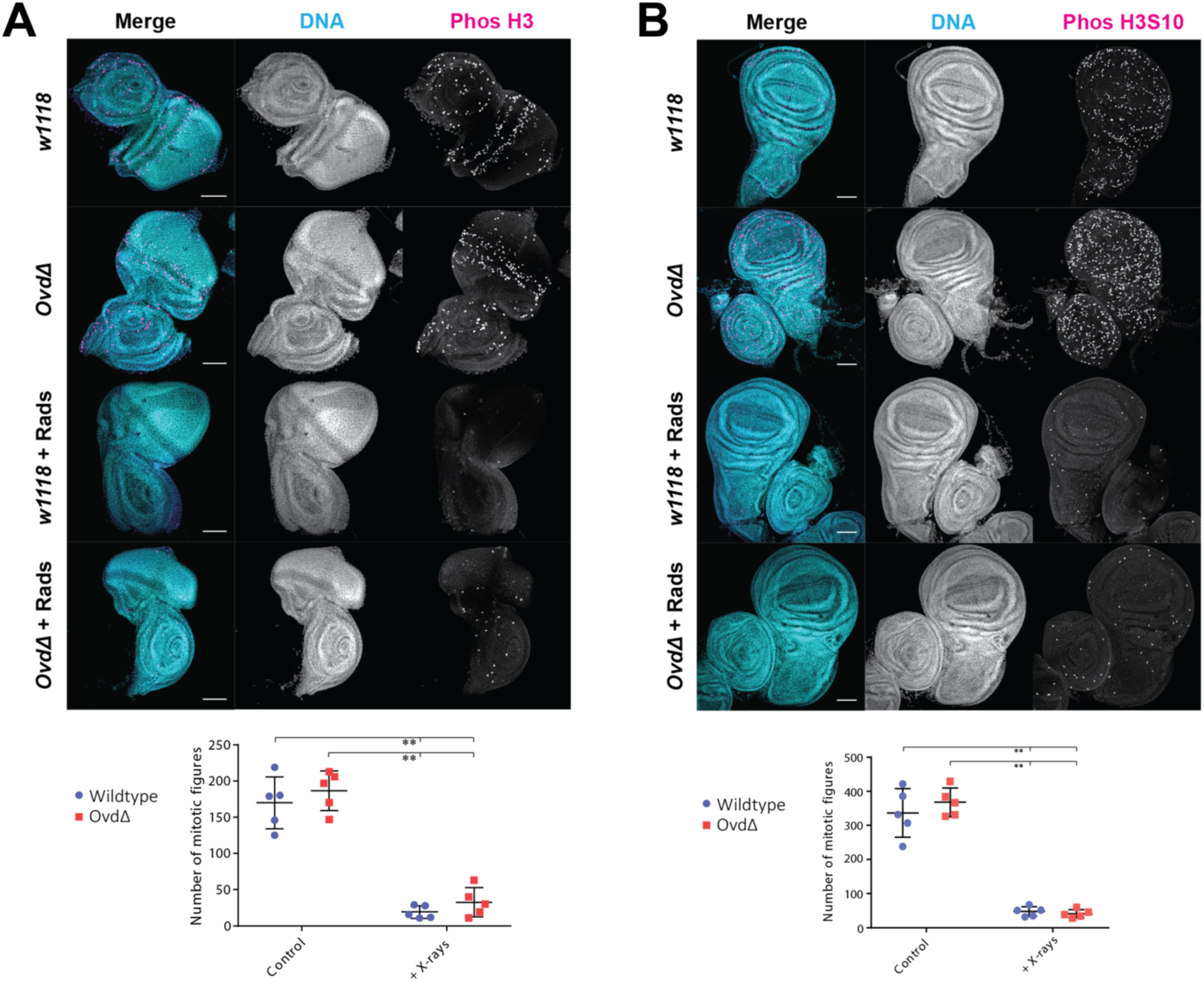
*Overdrive* knockout does not affect cell cycle progression in **(A)** eye discs or **(B)** wing discs. Discs from *OvdΔ* and *w^1118^* larvae were stained for DNA (cyan) and the mitotic marker H3S10Phos (magenta) after irradiation. Both *OvdΔ* and *w^1118^* largely arrest division after radiation, and the number of mitotic foci does not differ significantly between genotypes either before or after irradiation. Points represent biological replicates, and error bars represent standard deviation. *\*\* = p* < 0.005, Mann-Whitney test.

**Supplemental Figure S5:**
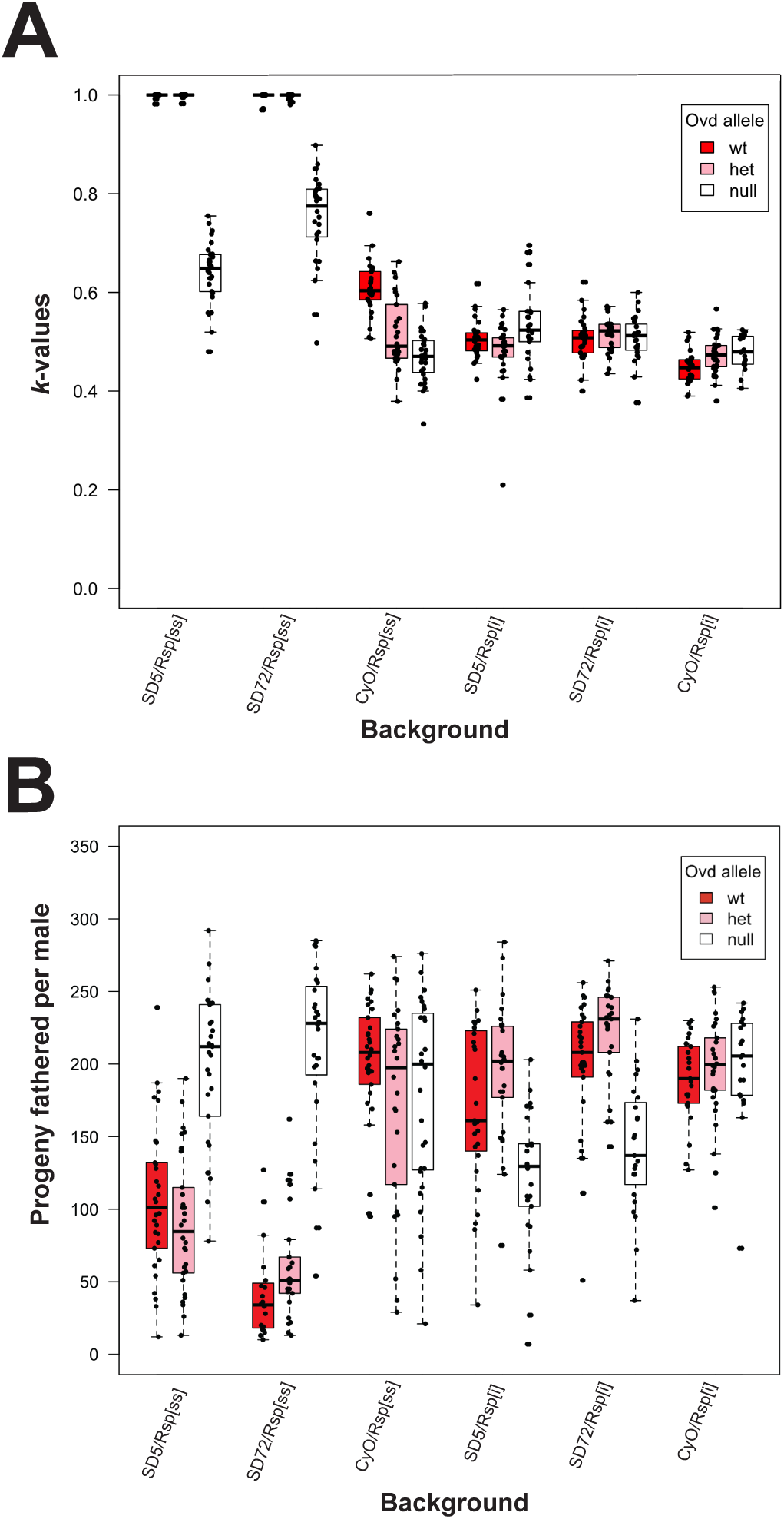
Expanded comparison of various *SD* and *Rsp* chromosomes in wild-type *Ovd*, heterozygous *Ovd^Δ^* and homozygous *Ovd^Δ^* backgrounds. **(A)** Proportion of progeny of tester males inheriting a *Segregation Distorter* chromosome (*SD5* or *SD72*) or non-driving *CyO* control. **(B)** Total number of offspring sired by each tester male. *Overdrive* knockout appears to restore fertility relative to *SD*-inactive controls in *SD/Rsp^ss^* backgrounds, even as it reduces fertility in *SD/Rsp^i^* backgrounds. For both panels, the center bar of boxes represents the median, and whiskers extend up to 1.5× the interquartile range.

## REFERENCES

1. L. Sandler, E. Novitski, Meiotic Drive as an Evolutionary Force. Am. Nat. 91, 105–110 (1957).

2. T. W. Lyttle, Segregation distorters. Annu. Rev. Genet. 25, 511–557 (1991).

3. C. Courret, C.-H. Chang, K. H.-C. Wei, C. Montchamp-Moreau, A. M. Larracuente, Meiotic drive mechanisms: lessons from Drosophila. Proc. Biol. Sci. 286, 20191430 (2019).

4. C. Merrill, L. Bayraktaroglu, A. Kusano, B. Ganetzky, Truncated RanGAP encoded by the Segregation Distorter locus of Drosophila. Science 283, 1742–1745 (1999).

5. Q. Helleu, P. R. Gérard, R. Dubruille, D. Ogereau, B. Prud’homme, B. Loppin, C. Montchamp-Moreau, Rapid evolution of a Y-chromosome heterochromatin protein underlies sex chromosome meiotic drive. Proceedings of the National Academy of Sciences 113, 4110–4115 (2016).

6. Y. Tao, L. Araripe, S. B. Kingan, Y. Ke, H. Xiao, D. L. Hartl, A sex-ratio meiotic drive system in Drosophila simulans. II: an X-linked distorter. PLoS Biol. 5, e293 (2007).

7. N. Phadnis, H. A. Orr, A single gene causes both male sterility and segregation distortion in Drosophila hybrids. Science 323, 376–379 (2009).

8. A. M. Larracuente, D. C. Presgraves, The selfish Segregation Distorter gene complex of Drosophila melanogaster. Genetics 192, 33–53 (2012).

9. D. Policansky, J. Ellison, “Sex Ratio” in Drosophila pseudoobscura: Spermiogenic Failure. Science 169, 888–889 (1970).

10. J. Vedanayagam, M. Herbette, H. Mudgett, C.-J. Lin, C.-M. Lai, C. McDonough-Goldstein, S. Dorus, B. Loppin, C. Meiklejohn, R. Dubruille, E. C. Lai, Essential and recurrent roles for hairpin RNAs in silencing de novo sex chromosome conflict in Drosophila simulans. PLoS Biol. 21, e3002136 (2023).

11. K. T. Tokuyasu, W. J. Peacock, R. W. Hardy, Dynamics of spermiogenesis in Drosophila melanogaster. I. Individualization process. Z. Zellforsch. Mikrosk. Anat. 124, 479–506 (1972).

12. M. T. Fuller, “Spermatogenesis” in The Development of Drosophila Melanogaster, A. A. M. M Bate, Ed. (Cold Spring Harbor Laboratory Press, New York, NY, 1993), pp. 71–148.

13. S. Jayaramaiah Raja, R. Renkawitz-Pohl, Replacement by Drosophila melanogaster protamines and Mst77F of histones during chromatin condensation in late spermatids and role of sesame in the removal of these proteins from the male pronucleus. Mol. Cell. Biol. 25, 6165–6177 (2005).

14. C. Rathke, W. M. Baarends, S. Jayaramaiah-Raja, M. Bartkuhn, R. Renkawitz, R. Renkawitz-Pohl, Transition from a nucleosome-based to a protamine-based chromatin configuration during spermiogenesis in Drosophila. J. Cell Sci. 120, 1689–1700 (2007).

15. L. Sandler, Y. Hiraizumi, I. Sandler, Meiotic Drive in Natural Populations of Drosophila Melanogaster. I. the Cytogenetic Basis of Segregation-Distortion. Genetics 44, 233–250 (1959).

16. M. Herbette, X. Wei, C.-H. Chang, A. M. Larracuente, B. Loppin, R. Dubruille, Distinct spermiogenic phenotypes underlie sperm elimination in the Segregation Distorter meiotic drive system. PLoS Genet. 17, e1009662 (2021).

17. C. I. Wu, T. W. Lyttle, M. L. Wu, G. F. Lin, Association between a satellite DNA sequence and the Responder of Segregation Distorter in D. melanogaster. Cell 54, 179–189 (1988).

18. E. Hauschteck-Jungen, D. L. Hartl, Defective Histone Transition during Spermiogenesis in Heterozygous *Segregation Distorter* Males of *Drosophila melanogaster*. Genetics 101, 57– 69 (1982).

19. B. T. Wakimoto, D. L. Lindsley, C. Herrera, Toward a comprehensive genetic analysis of male fertility in Drosophila melanogaster. Genetics 167, 207–216 (2004).

20. G. Ben-David, E. Miller, J. Steinhauer, Drosophila spermatid individualization is sensitive to temperature and fatty acid metabolism. Spermatogenesis 5, e1006089 (2015).

21. J. Steinhauer, Separating from the pack: Molecular mechanisms of Drosophila spermatid individualization. Spermatogenesis 5, e1041345 (2015).

22. B. D. McKee, K. Wilhelm, C. Merrill, X. Ren, Male sterility and meiotic drive associated with sex chromosome rearrangements in Drosophila. Role of X-Y pairing. Genetics 149, 143– 155 (1998).

23. L. D. Hurst, A. Pomiankowski, Causes of sex ratio bias may account for unisexual sterility in hybrids: a new explanation of Haldane’s rule and related phenomena. Genetics 128, 841– 858 (1991).

24. S. A. Frank, Divergence of meiotic drive-suppression systems as an explanation for sex-biased hybrid sterility and inviability. Evolution 45, 262–267 (1991).

25. M. M. Patten, Selfish X chromosomes and speciation. Mol. Ecol. 27, 3772–3782 (2018).

26. J. A. Coyne & H. A. Orr, Speciation (Sinauer Associates, Sunderland, MA, 2004).

27. J. Wen, H. Duan, F. Bejarano, K. Okamura, L. Fabian, J. A. Brill, D. Bortolamiol-Becet, R. Martin, J. G. Ruby, E. C. Lai, Adaptive regulation of testis gene expression and control of male fertility by the Drosophila hairpin RNA pathway. [Corrected]. Mol. Cell 57, 165–178 (2015).

28. H. A. Orr, S. Irving, Complex epistasis and the genetic basis of hybrid sterility in the Drosophila pseudoobscura Bogota-USA hybridization. Genetics 158, 1089–1100 (2001).

29. H. A. Orr, S. Irving, Segregation distortion in hybrids between the Bogota and USA subspecies of Drosophila pseudoobscura. Genetics 169, 671–682 (2005).

30. N. Phadnis, Genetic Architecture of Male Sterility and Segregation Distortion in Drosophila pseudoobscura Bogota–USA Hybrids. Genetics 189, 1001–1009 (2011).

31. A. C. M. Junqueira, A. M. L. Azeredo-Espin, D. F. Paulo, M. A. T. Marinho, L. P. Tomsho, D. I. Drautz-Moses, R. W. Purbojati, A. Ratan, S. C. Schuster, Large-scale mitogenomics enables insights into Schizophora (Diptera) radiation and population diversity. Sci. Rep. 6, 1–13 (2016).

32. B. M. Wiegmann, M. D. Trautwein, I. S. Winkler, N. B. Barr, J.-W. Kim, C. Lambkin, M. A. Bertone, B. K. Cassel, K. M. Bayless, A. M. Heimberg, B. M. Wheeler, K. J. Peterson, T. Pape, B. J. Sinclair, J. H. Skevington, V. Blagoderov, J. Caravas, S. N. Kutty, U. Schmidt-Ott, G. E. Kampmeier, F. C. Thompson, D. A. Grimaldi, A. T. Beckenbach, G. W. Courtney, M. Friedrich, R. Meier, D. K. Yeates, Episodic radiations in the fly tree of life. Proc. Natl. Acad. Sci. U. S. A. 108, 5690–5695 (2011).

33. H. A. Orr, J. P. Masly, N. Phadnis, Speciation in Drosophila: from phenotypes to molecules. J. Hered. 98, 103–110 (2007).

34. P. L. Oliver, L. Goodstadt, J. J. Bayes, Z. Birtle, K. C. Roach, N. Phadnis, S. A. Beatson, G. Lunter, H. S. Malik, C. P. Ponting, Accelerated evolution of the Prdm9 speciation gene across diverse metazoan taxa. PLoS Genet. 5, e1000753 (2009).

35. Z. Yang, PAML 4: phylogenetic analysis by maximum likelihood. Mol. Biol. Evol. 24, 1586– 1591 (2007).

36. J. H. Lee, E. Lee, J. Park, E. Kim, J. Kim, J. Chung, In vivo p53 function is indispensable for DNA damage-induced apoptotic signaling in Drosophila. FEBS Lett. 550, 5–10 (2003).

37. J. Xu, S. Xin, W. Du, Drosophila Chk2 is required for DNA damage-mediated cell cycle arrest and apoptosis. FEBS Lett. 508, 394–398 (2001).

38. G. Cutler, K. M. Perry, R. Tjian, Adf-1 is a nonmodular transcription factor that contains a TAF-binding Myb-like motif. Mol. Cell. Biol. 18, 2252–2261 (1998).

39. X. Yi, H. I. de Vries, K. Siudeja, A. Rana, W. Lemstra, J. F. Brunsting, R. M. Kok, Y. M. Smulders, M. Schaefer, F. Dijk, Y. Shang, B. J. L. Eggen, H. H. Kampinga, O. C. M. Sibon, Stwl modifies chromatin compaction and is required to maintain DNA integrity in the presence of perturbed DNA replication. Mol. Biol. Cell 20, 983–994 (2009).

40. D. Zinshteyn, D. A. Barbash, Stonewall prevents expression of ectopic genes in the ovary and accumulates at insulator elements in D. melanogaster. PLoS Genet. 18, e1010110 (2022).

41. R. L. Kurzhals, S. W. A. Titen, H. B. Xie, K. G. Golic, Chk2 and p53 are haploinsufficient with dependent and independent functions to eliminate cells after telomere loss. PLoS genetics 7, e1002103 (2011).

42. S. J. Gratz, F. P. Ukken, C. D. Rubinstein, G. Thiede, L. K. Donohue, A. M. Cummings, K. M. O’Connor-Giles, Highly specific and efficient CRISPR/Cas9-catalyzed homology-directed repair in Drosophila. Genetics 196, 961–971 (2014).

43. T. Dunckley, M. Tucker, R. Parker, The teflon gene is required for maintenance of autosomal homolog pairing at meiosis I in male Drosophila melanogaster. Genetics 157, 273–281 (2001).

44. 44. J. Schindelin, I. Arganda-Carreras, E. Frise, Fiji: an open-source platform for biological-image analysis. Nature Methods 9, 676–682 (2012).

45. R. C. Edgar, MUSCLE: multiple sequence alignment with high accuracy and high throughput. Nucleic Acids Research 32, 1792–1797 (2004).

46. 46. A. M. Waterhouse, J. B. Procter, D. M. A. Martin, M. Clamp, G. J. Barton, Jalview Version 2 – a multiple sequence alignment editor and analysis workbench. Bioinformatics 25, 1189–1191 (2009).

47. A. Drozdetskiy, C. Cole, J. Procter, G. J. Barton, JPred4: a protein secondary structure prediction server. Nucleic Acids Research 43, W389–W394 (2015).

48. J. C. Cooper, N. Phadnis, Parallel Evolution of Sperm Hyper-Activation Ca2+ Channels. Genome Biology and Evolution 9, 1938–1949 (2017).

49. L. Sandler, Y. Hiraizumi, Meiotic drive in natural populations of *Drosophila melanogaster*. ii. genetic variation at the segregation-distorter locus. Proceedings of the National Academy of Sciences 45, 1412–1422 (1959).

